# Sub-Confluent Culture of Mouse Embryonic Stem Cell-derived Ventricular Cardiomyocytes On & In Gels - Enhancement of Maturation Phenotype Relative to Tissue Culture Polystyrene via Enabling of Auxotonic Contraction

**DOI:** 10.1101/199737

**Authors:** Nikhil Mittal, Ayhan Atmanli, Dongjian Hu, Daniel Groeneweg, Eduardo Kausel, Ibrahim J Domian

## Abstract

Cardiac myocytes (CMs) obtained by differentiating embryonic stem cells (ES-CMs) have an immature phenotype and promoting the maturation of such PSC-derived cardiomyocytes remains a major limitation in the development of stem cell models of human cardiovascular disease. We cultured murine ES-CMs in a collagen gel (3D) at a low density, or on collagen-coated polystyrene (2D) and found that 3D culture results in dramatic improvement of the maturation rate and end-state gene expression of ES-CMs. There are two main differences between CMs cultured in 3D versus 2D; in 3D the mechanical stiffness of the environment is lower, enabling auxotonic instead of isometric contraction; and, in 3D the amount of cell-cell interaction is higher. To isolate the contributions, we first cultured ES-CMs *on* gels (2D substrates) that are softer than tissue culture plastic, enabling auxotonic contraction, while controlling for dimensionality and cell interaction. This indeed promoted a mature gene expression profile, while also enabling the maintenance of sarcomeres. Next, we determined that increased cell-cell interaction inhibits the mature gene expression of ES-CMs. Thus, auxotonic contraction is the likely mechanism for improved gene expression in sub-confluent 3D culture. However, 2D auxotonic contraction may offer a suitable compromise between obtaining enhanced gene expression and morphology. After 6 weeks of culture on gels, via Di-8-ANEPPs and WGA staining we also detected CMs forming a t-tubule network. Collectively these results demonstrate that 3D and 2D cultures that enable auxotonic contraction enhance aspects of the maturation of ES-CMs.

## Introduction

Pluripotent stem cells (PSCs) are a renewable source for obtaining any kind of cell present in the adult organism. Indeed, PSCs offer tremendous promise for the generation of patient specific cells for drug screening, cell therapy, and the in vitro generation of transplantable tissue. Over the last decade the focus of the field has been on developing protocols for the directed differentiation of PSCs into specific cell types so as to be able to generate large numbers of the cell of interest, and this effort has largely met with success. Cells thus obtained could be, and to some extent are, used for a multitude of applications – from studying their biology, to testing therapies against disease. However, this latter application, especially, suffers from the problem that the cells obtained by differentiating PSCs typically retain an immature neonatal-like phenotype. While work from several laboratories has demonstrated the ability to differentiate PSCs into progenitor cells of various types, driving these cells toward a more mature adult-like phenotype remains a major challenge in the field. In fact, mature cells only appear to have been demonstrated for neuronal cells such as motor neurons [1] and dopamine neurons [2, 3]; and erythrocytes [4]. For most cell types, including beta cells [5], hepatocytes [6], and cardiomyocytes (reviewed in [7, 8]), differentiated cells obtained from PSCs show a largely immature phenotype.

Almost all studies to date on the maturation of embryonic stem cell (ESC)-derived cardiomyocytes (hereafter referred to as ES-CMs) have focused on human ES-CMs cultured in a “traditional” manner i.e. on tissue culture polystyrene (TCPS). With respect to their electrophysiological characteristics, after ~2 months in culture ES-CMs express some potassium current-associated proteins at levels similar to that found in the adult human heart [9, 10]. Additionally, a recent study found adequate binucleation after 100 days in culture [11]. However, even after 1-2 months in culture, transverse tubules (t-tubules) could not be detected in ES-CMs [12, 13]. Lundy *et al.* did not observe t-tubules even after 4 months in culture [11]. Further, Liu *et al.* [14] found that they did not express calcium-handling proteins such as calsequestrin, triadin, and junctin (after ~3 weeks in culture), which are necessary for calcium-induced-calcium release. Similarly, Dolnikov *et al.* [15] also found that ES-CMs did not express phospholamban or calsequestrin (7-55 days). Finally, ES-CMs had limited (physiological) hypertrophy at 3 months [12].

A parallel stream of research is cardiac tissue engineering which concerns itself with developing adult heart-like tissue constructs *in vitro*. This is achieved by suspending the cells in a matrix usually made up of extracellular matrix (ECM) proteins. Additionally, the cells are allowed to achieve confluence via plating at high densities. Much of the early work in this field was done using neonatal rat CMs, but some recent studies have used ES-CMs [16, 17]. After 2 weeks of culture in a fibrin-matrigel scaffold with isometric contraction imposed (starting at day 22-30 of differentiation), Zhang *et al.* [16] have recently reported that they obtained stresses that were 60% of the adult human wall stress, similar to neonatal wall stress [18]. After 7 days of culture in a collagen scaffold with isometric contraction imposed (starting at day 20 of differentiation), another study recently reported that they obtained conduction velocities similar to that of adult human heart tissue [17]. Turnbull *et al.* similarly cultured ES-CMs in a collagen gel suspended between silicone posts for 18-24 days (starting at day 15 of differentiation) and determined that the resulting constructs exert forces that approach those in the neonatal myocardium [19]. Gene expression also differed significantly from the adult heart. Schaaf *et al*. showed that similarly cultured ES-CMs are also electrophyiologically immature [20]. Studies have also shown that electrical stimulation leads to maturation of cells within such constructs [21, 22], though the cells did not achieve adult-like maturity. In contrast culturing *human* ES-CMs on soft substrates appeared to result in no improvement in force generation after 1 month of culture [23]. Collectively these results suggest that 3D culture may be more effective at promoting ES-CM maturation relative to (2D) culture on polystyrene culture dishes. In this study, we wanted to examine this hypothesis further, and investigate the underlying mechanisms.

The primary differences between culture in gels versus on polystyrene are (i) that the gels are soft, enabling auxotonic contraction (i.e. resistance increases with contraction length) especially at sub-confluent densities (i.e. if the matrix is not digested away by the cells), and (ii) the cell-cell interaction may be higher in three dimensions due to the additional interactions of cells with the cells above and below them. For neonatal rat ventricular CMs, it has been shown that auxotonic contraction improves maturation relative to isometric contraction [24]. In contrast culturing *human* ES-CMs on soft substrates appeared to result in no improvement in force generation after 1 month of culture [23]. This raises an additional question as to whether this difference between the result for neonatal rat CMs versus human ES-CMs has to do with the developmental stage of the original cells i.e. neonatal versus embryonic, or to do with the species. Our results with mouse ES-CMs demonstrate that auxotonic contraction does enhance maturation even for ES-derived CMs. Thus, the above difference is likely due to species-specific effects.

## Results

We validated a panel of 8 genes that are implicated in both major aspects of CM function, namely, force generation (MHCs), and calcium-induced calcium release (Ryr2, Casq2, Trdn, Serca2a); as well as Calr and Cx40 (Figure 1a). The full names of these genes and the motivation for including them in our panel are described in detail in the supplementary information. We chose these genes in part based on previous studies that have shown several of them to be also associated with the physiological maturation of human CMs [14]. We favored these genes over ion channels usually studied in this field i.e. typically ion channels implicated in drug-induced Torsades de Pointes because there are large differences in the roles of various ion currents in mouse versus human CMs [25].

We had previously developed a transgenic system for isolating Nkx2.5+AHF+ (immature) cells from mice as well as mouse ESCs [26]. These cells are highly enriched for early/immature ventricular CMs (> 80%, EVCMs), enabling the study of maturation of CMs with minimal confounding effects related to differences in the maturation of nodal versus atrial versus ventricular cells, or the commitment of other cell types to the cardiac lineage.

We cultured these EVCMs for 2 weeks either on tissue culture polystyrene (TCPS) coated with collagen I (200 μg/ml, 30 minutes), or within a collagen I gel (200 μg/ml) (Figure 1b). On polystyrene as well as in the gel, spontaneous beating activity was observed (Supplementary Movies S1 and S2). We found that culture within the gel led to the formation of 3D colonies. Importantly, this 3D culture led to a dramatic improvement in the expression of most genes in our panel (Figure 1c). Expression of calreticulin (Calr) was lower in 3D, consistent with the fact that Calr expression negatively correlates with maturation (Figure 1a). Additionally, expression of MHCB as well as Cx40 were below the detectable limit for the cells cultured in the gel, and highly variable on polystyrene, perhaps due to the low level of expression.

The cells plated on polystyrene were 90 ± 6% troponin-positive indicating that this difference in gene expression was not due to the overgrowth of other cell types. This can also be appreciated via the alpha-actinin immunostaining performed (Figure 1d).

We next measured the kinetics of gene expression changes for a few key genes. In particular, we wanted to answer the following question - in 2D do the genes never get up-regulated? Or do they go up but subsequently decline? We measured the variation with time of the expression of MHCA (as a representative gene involved in mechanical function) and Ryr2 (as a representative gene involved in electrical function) relative to adult left ventricular (LV) tissue (Figure 1e). We found that culture on TCPS alone led to a minimal increase in the expression of these genes after 10 days of culture, and culture for an additional 4 days led to no further increase in the expression of these genes. In contrast to this data, culture in 3D led to a rapid increase in the expression of these genes, reaching adult-like expression levels in only 10 days. Correspondingly, for Calreticulin, for which expression correlates negatively with maturation, expression declined rapidly in 3D to adult-like values, but remained high on polystyrene (Figure 1f). These data demonstrate that mature gene expression is not achieved for culture on polystyrene at any time during the 2-week period.

These results demonstrating that the sub-confluent/low density 3D culture of ES-CMs can result in adult-like gene expression are novel. However, in terms of morphology, the cells did not adopt the rod-like morphology of mature cardiomyocytes (Figure 1g), perhaps because we were unable to stretch the gels (see Discussion). Additionally, the expression of some genes was super-physiological (Figure 1c), which may again be due to the lack of a resistance/afterload (Discussion).

### Improved Gene Expression in 3D: Auxotonic contraction versus Isometric contraction

We then aimed to further understand why 3D culture results in enhanced gene expression. There are two main differences between the cells cultured within the gel versus on TCPS (Figure 2a). (i) The cells can shorten in 3D, since they are not bound to a hard surface (TCPS), and (ii) cell-cell interaction may be greater in 3D since cells can additionally interact with cells above and below themselves.

**Figure 1:**
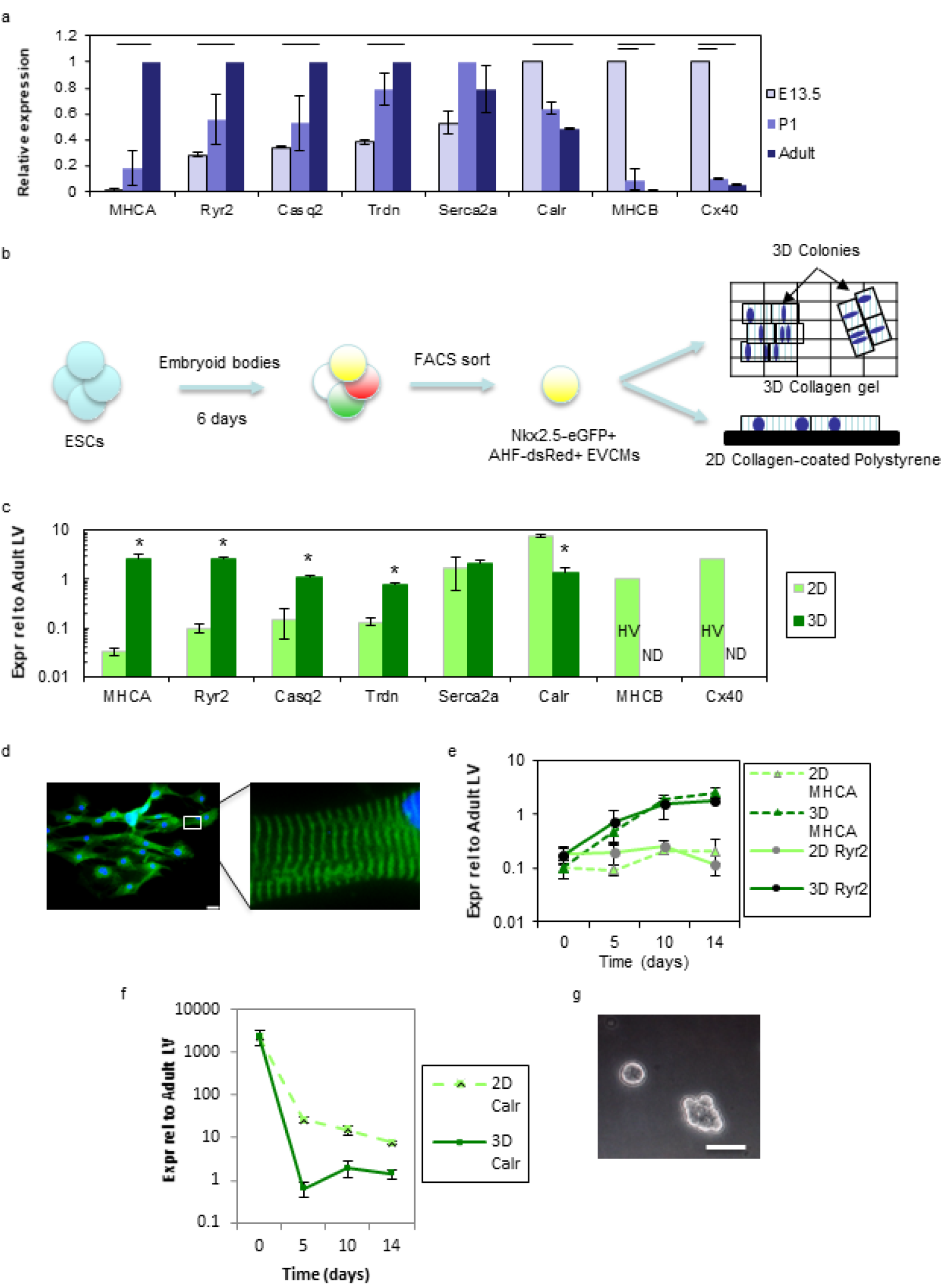
The effect of cellular organization on cardiac maturation. (a) Relative expression for several genes, for left ventricles obtained from mouse E13.5 embryos, postnatal day 1 (PD1), and adult mice. (b) Schematic of experimental design. ESCs were cultured as embryoid bodies for 6 days, followed by dissociation and FACS to obtain early ventricular cardiac myocytes (EVCMs) as previously described. (c) Relative expression for several genes for cells plated on polystyrene coated with collagen I (2D), or in a collagen gel (3D), for 2 weeks. (d) Fluorescence images of EVCMs plated on collagen-coated polystyrene for 2 weeks. Stained with α-actinin antibody (green) and DAPI (blue). (e) Time course of the expression of MHCA (triangles) and Ryr2 (circles) for cells grown in 2D (dashed lines) and 3D (solid lines). (f) Time course of the expression of Calr for cells grown in 2D (dashed line) and 3D (solid line). (g) Phase contrast image of clusters formed by EVCMs plated in a collagen gel for 2 weeks. Scale bars represent 50μm. Error bars represent SD. * and lines represent p < 0.05. HV: Highly variable; ND: Not detectable

We first investigated the effect of EVCM auxotonic contraction on maturation. On polystyrene (Young’s modulus = 3 GPa) even an adult CM would only be able to provide sufficient force to undergo a shortening of approximately 0.00005% (Supplementary Information). As a result, contracting cardiomyocytes effectively experience isometric contraction. Experimentally, we often observed that the central regions of colonies detached from the substrate (TCPS), leaving only the colony boundary attached to the substrate. In such colonies, we observed that while cells in the central region of the cluster were able to shorten, cells towards the periphery were forced to undergo compensatory lengthening during systole (Supplementary Movie S2). This latter phenomenon is highly unphysiological, and could explain the overall lack of maturation generally observed on TCPS and other systems that impose isometric contraction.

Previous studies have shown that CMs <u>from other (non-ES) sources</u> can be cultured *on* (instead of *in*) soft materials (typically polyacrylamide (PAAm)), enabling auxotonic contraction [23, 27, 28]. Indeed, culture of neonatal rat ventricular myocytes [28] and embryonic chicken myocytes [29] on such substrates leads to their partial maturation. To validate our hypothesis, we therefore decided to use this strategy to test the effect of auxotonic versus isometric contraction on the maturation of <u>ES-derived ventricular CMs</u>.

Culture on these substrates indeed enabled auxotonic contraction (Figure 2b, Supplementary Movie S3). <u>The amount of fractional shortening was inversely related to the resistance i.e. substrate stiffness, and it increased with time in culture consistent with functional maturation of the cells</u> (Figure 2c). Importantly, culture on these substrates for 2 weeks resulted in more adult-like gene expression relative to culture on collagen coated TCPS (Figure 2d). These results demonstrate that auxotonic contraction enables mature gene expression in ES-derived CMs.

Next, we examined cell-cell interactions. In 2D, cells can only interact with their neighbors in the same plane, while in 3D they can additionally interact with cells above and below themselves and therefore experience higher cell-cell interaction (paracrine *and* juxtacrine). To vary the total amount of cell-cell interaction (paracrine and juxtacrine), while controlling for dimensionality, we either plated cells at varying densities on polystyrene, or made 3D clusters with varying numbers of cells. We found that *decreasing* the 2D plating density (by 100-fold) actually led to *improved* gene expression (Figure 2e). In 3D it is not possible to achieve such a high contrast in cell number because in large clusters the central region becomes necrotic due to oxygen and nutrient deprivation. We created clusters with either 100 or 500 EVCMs and did not detect significant differences in gene expression (Figure 2f). These results indicate that increased cell-cell interaction is not the primary mechanism that leads to mature gene expression in 3D culture. While we cannot completely rule out differences in the depletion of culture components as a potential cause for the observed effects, the cell densities were quite low (15000 cells/ml), and we performed half-medium changes every other day. Thus, it is unlikely that nutrient depletion or metabolite accumulation played a significant role. Previous studies demonstrated that these effects generally occur at cell densities of around 500,000 cells/ml and 3-4 days without feeding [30]. Additionally, we are not aware of any other method that can simultaneously modulate the total paracrine and juxtacrine signal.

Together these results demonstrate that 3D culture enhances the maturation of mouse ES-derived EVCMs by enabling auxotonic contraction.

### Further Characterization of Mouse ES-CMs Undergoing 2D Auxotonic Contractions

Our initial results (Figure 2b-d) suggested that culture on 2D gels may provide a suitable compromise between obtaining rapid improvement in gene expression versus obtaining cells with a more elongated morphology. Repeated auxotonic contractions constitute dynamic resistance training (DRT). So we wanted to further characterize the effect of such 2D DRT on some of the functional and morphological properties of ES-CMs. We have previously demonstrated that the EVCMs used in this study have ventricular action potentials [26]. Next, we used traction force microscopy (methods) to determine the forces exerted by trained ES-CMs, and found that force increased with increasing resistance i.e. substrate stiffness (Figure 2g). Similar results have been previously observed for neonatal rat CMs [27]. <u>In a previous essay, we have discussed in detail why this stiffness value may differ from the typical stiffness of the heart wall</u> [31]. For gels with a modulus of 50 and 100 kPa, the average force was 2.5–3 μN/cell (at 2 weeks), which is in the range known to be exerted by adult CMs in vivo (see below) for mice from the C57BL/6 strain, which is the same strain from which our ES line was derived. To determine this latter value, we obtained the product of the wall stress (91 g/cm^2^=9 kPa [32]) with the cross-sectional area (284–568 μm^2^ [33]). Our values are also within the range obtained for rat CMs using traction force microscopy [27, 28].

**Figure 2.**
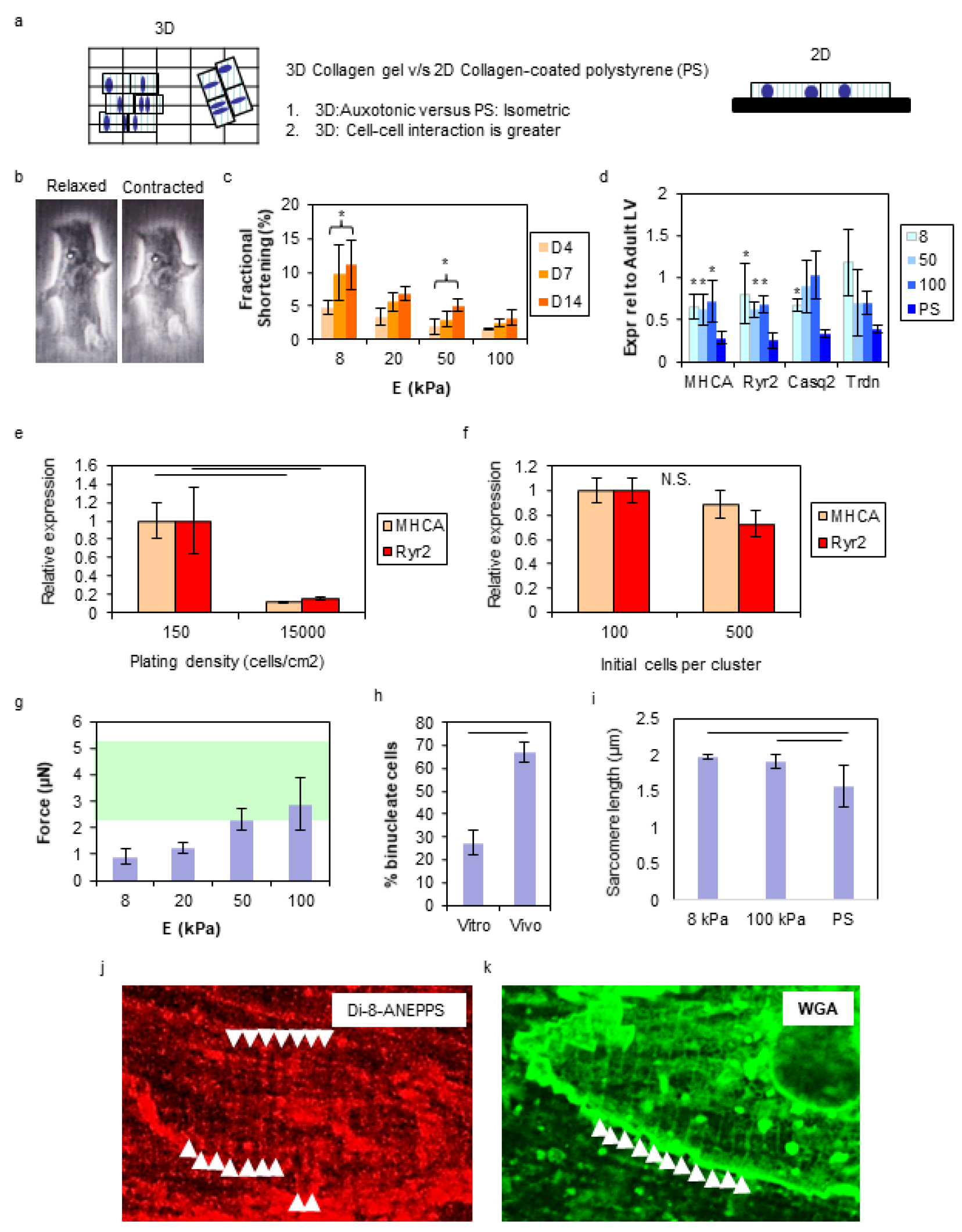
The effect of auxotonic contraction on mouse ES-CM maturation. (a) Schematic showing the differences between culture in a gel and on polystyrene. (b) An EVCM plated on a 8 kPa PAAM gel for 7 days undergoes fractional shortening when it beats. (c) Fractional shortening of EVCMs plated on gels of various stiffnesses on day 4, day 7, and day 14 of culture. (d) Gene expression relative to adult LV for cells plated on TCPS or PAAm. (e) Gene expression for EVCMs plated on polystyrene at two densities for two weeks. (f) Gene expression for EVCMs in clusters of 100 or 500 cells for two weeks. (g) Force generated by EVCMs cultured on gels for two weeks. The green region represents the range for adult CM. (h) Percent bi-nucleate cells for EVCM on 100 kPa gels (Vitro) and adult CM (Vivo). (i) Sarcomere length at 6 weeks. (j) Confocal image of EVCM cultured on a 100 kPa gel for 6 weeks and treated with Di-8-ANEPPS. Arrowheads point to regions of linear staining. (k) Confocal image of EVCM cultured on a 100 kPa gel for 6 weeks and stained with wheat germ agglutinin (WGA). Arrowheads point to regions of linear staining. Error bars represent SD. * and lines represent p < 0.05

In a previous study, we demonstrated that EVCMs exit the cell cycle after 4–5 days in culture [26]. Thus, cells with two nuclei at 14 days are terminally binucleate cells. We determined the percentage of binucleate cells (2 weeks, 100 kPa gels). Binucleation was considerably lower than that observed in vivo in CMs extracted from ventricles of 12-week animals from the same transgenic line that the ES cells were derived from (Figure 2h). However, this is not an impediment to downstream applications, since the cells can be sorted and reconstituted at the appropriate ratio. CM maturation is also associated with an increase in sarcomere length [34, 35]. Sarcomere lengths in trained ES-CMs (6 weeks), were comparable to the typical sarcomere lengths of adult mouse CMs described in the literature [36, 37] (Figure 2i). Meanwhile, similar to results reported for neonatal rat CMs, after 6 weeks of culture on PS, staining for α-actinin was very weak, consistent with loss of sarcomere integrity [38]. Finally, we assayed trained ES-CMs for the presence of t-tubules. For Di-8-ANEPPS [39, 40] which only works on live cells we had to keep the gel hydrated during the imaging process by placing it in a glass bottom dish. Additionally, due to the need to image through the hydrogel (and backing coverslip), we were maximally able to use a 40x lens. For CMs that had undergone 2 or 4 weeks of DRT we did not detect t-tubules. However, for CMs that underwent 6 weeks of DRT, we detected CMs with a mature staining pattern in regions of the cell (Figure 2j). This was further confirmed with staining with the membrane stain wheat germ agglutinin (WGA) (Figure 2k).

## Discussion

The long-awaited applications of PSCs in developing new therapies for heart disease require the ability to generate adult-like CMs from PSCs. In this study, we have taken advantage of a previously developed source of ESC-derived murine ventricular CMs [26] to study the effects of 3D culture on ES-CM maturation. We believe that this is the first study to do a side-by-side comparison of culture in gels versus polystyrene for ES-CMs.

While the current cell source i.e. Nkx2.5+AHF+ cells are relatively stable cell source in terms of the fraction of myocytes generated etc., these cells are relatively rare (within embryoid bodies). Thus, in this study we typically obtained low numbers of these cells and consequently plated them at low, sub-confluent cell densities. In future work it would be interesting to optimize the cell density by using an alternate more abundant cell source or developing protocols to obtain larger quantities of Nkx2.5+AHF+ cells.

Tissue culture polystyrene (TCPS) is the substrate of choice for culturing most cells. However, as it is a stiff material, colonies of CMs grown on it are unable to undergo net shortening. Thus, while some CMs undergo shortening, other CMs in the colony must undergo a compensatory lengthening during systole, which is highly unphysiological (Supplementary Movie S2). We found that culture on TCPS alone led to a minimal increase in the expression of maturation genes after 10 days of culture, and culture for an additional 4 days led to no further increase in the expression of these genes, suggesting that merely extending the culture period would not lead to the maturation of cells growing on TCPS (Figures 1e and 1f). In contrast to this data, culture in 3D led to a rapid improvement in the expression of these genes, reaching adultlike expression levels in only 10 days. However, since the CM maturation period in mice is ~30 days [41], we cannot rule out that longer durations of 2D culture would lead to maturation. Additionally, while the gene expression did not increase in the day 10 – day 14 period, it is possible that protein expression/organization increased. However, a previous study has shown that a fraction of mouse ES-CMs cultured on TCPS for 4-5 weeks undergo sarcomeric disassembly [38]. Thus, for this application, TCPS is an unsuitable culture substrate. In other studies, tissue constructs have been made by embedding ES-CMs into fibrin or collagen gels at high densities [16, 17]. In these systems again, the contraction at least at the construct level is isometric and thus the phenomenon described above may occur. For neonatal rat ventricular cardiomyocytes, Hirt *et al.* have also shown that for such constructs, isometric contraction leads to a heart failure phenotype [42]. Additionally, in tissue constructs, since the cells themselves compact the gel [16, 17, 19, 20], there is no direct control over the construct stiffness i.e. the resistance the cells work against. As we have observed, the value of this resistance strongly impacts the force generated by these cells i.e. their functional maturation (Figure 2g), and thus it is likely important to control this parameter.

We investigated the mechanism behind the maturation caused by 3D culture relative to 2D culture. We propose that are two differences between the culture types that may lead to an effect: the ability to undergo fractional shortening, and cell-cell interactions. Our 2D experiments on gels demonstrate that fractional shortening indeed improves gene expression. However, our results also indicate that shortening alone is insufficient/sub-optimal unless performed against an optimal resistance. In particular, the substrate on which the cells underwent the most fractional shortening (8 kPa condition) did not reach gene expression levels found in the 3D culture condition and produced the lowest traction forces. Similarly, in our unstretched 3D constructs, the CMs likely experience little or no resistance, and hence though they mature w.r.t. gene expression, they remain morphologically highly immature (Fig 1c). In this regard, a limitation of our study was the inability to stretch our 3D constructs. Previous studies have shown that mechanical stretching leads to CMs with an elongated morphology within 3D constructs [16, 20, 21]. Likely due to the lack of non-myocytes in our gels we did not observe compaction [43], which precluded us from stretching them. In future studies it would be interesting to mix ES-CMs with cardiac fibroblasts which may enable compaction and stretching of the gels. Finally, we observed that increasing cell-cell interactions actually impedes maturation (2D), or does not result in improved gene expression (3D), and therefore ruled it out as a contributing mechanism.

In a previous study [23] DRT did not lead to (statistically significant) time-dependent increase in force production in hESC-CM at one month. While we were able to observe increases in the force production by mouse ES-CMs within approximately one week, it is expected that this process may take longer in human ES-CM corresponding with the longer gestation period and lifespan in humans. In addition to species-specific differences, other factors could also contribute to the difference observed relative to our study. A mixture of atrial, ventricular and nodal CMs was used in that study. Nodal CMs are not involved in force generation and thus may not respond to DRT. Thus, the presence of a significant fraction of nodal cells could explain the lack of increase in force production observed. Also, in that study the gels were coated with fibronectin while our gels were coated with collagen I. Prior work from our lab [44] has demonstrated that ECM composition directly impacts myocardial contractility.

Finally, it is worth noting that in humans the CM maturation period has been described to take ~4 years [18], indicating the challenge associated with developing mature human CMs. Thus, while mimicking the in vivo environment may eventually lead to maturation, the monetary cost associated with such an approach will need to be considered. Inevitably pharmaceutical companies (and others) will need to weigh the cost versus benefits, and approaches that use other/hybrid approaches to obtain cells with limited maturity more rapidly and with substantially lower cost may eventually prove to be optimal. 3D culture or culture on soft substrates could be coupled with other approaches for driving the maturation of CMs such as genetic manipulation or addition of chemical inhibitors/activators. We anticipate that the ability to easily generate large quantities of mature CMs will be greatly beneficial in advancing the long-awaited applications of stem cells in developing new therapies for heart diseases.

## Materials and Methods

All animal procedures were performed in accordance with the NIH Guide for the Care and Use of Laboratory Animals and approved by the Institutional Animal Care and Use Committee at Massachusetts General Hospital.

### Mouse ES Culture

Murine Nkx2.5-eGFP and Anterior-heart-field-Mef2C-DsRed double transgenic embryonic stem (ES) cells were maintained on irradiated mouse embryonic fibroblasts in serum containing ESC medium. Prior to differentiation, cells were adapted for 2 days on 0.1% gelatin-coated polystyrene plates in adaptation medium. At day 0, cells were dissociated with 0.25% trypsin-EDTA for 2 minutes and re-suspended at a density of 100,000 cells/mL in differentiation medium. Differentiation was induced in embryoid bodies (EBs). On day 6 EBs were trypsinized to generate a single cell suspension and GFP+/DsRed+ EVCMs were isolated using FACS. We obtained sorting purities of >85%. Cells were subsequently cultured in differentiation medium, which was refreshed (50%) every other day.

### Polyacrylamide substrates

Polyacrylamide substrates of varying stiffnesses were prepared according to established protocols [27, 28]. Briefly, the stiffness of these PAAm substrates can be varied by varying the monomer and cross-linker concentrations [28]. We covalently linked collagen I to the PAAm using a heterobifunctional linker (Sulfo-SANPAH [27, 28]). Previous work has shown that the surface density of collagen I obtained by using this method of linkage is independent of substrate stiffness [27]. The stiffness values of the gels were obtained from Jacot *et al.* [28, 45].

### Immunocytochemistry

Cells were fixed in 4% paraformaldehyde for 7.5 minutes and stored in phosphate buffered saline at 4 °C. The α-actinin antibody (Sigma A7811) used was diluted 1:400 in blocking buffer. WGA conjugated with Alexa Fluor 488 was purchased from Thermo Scientific. <u>Sarcomere lengths</u> were determined using images of cells stained for α-actinin. The length of groups of 5-10 sarcomeres was measured using ImageJ (NIH). A total of 9 cells were measured from 2 batches of differentiation. <u>Percent binucleation</u> was determined by only using cells that had clear boundaries, typically isolated cells that were not a part of a colony. Data are from 3 biological replicates (9/34, 4/14, 15/45 cells).

### Collagen gels

20 μl of rat tail collagen I (3.5 mg/ml, BD Biosciences) was mixed with 20 μl of 0.1M NaOH. To this mixture we added 40 μl of fetal bovine serum. The identical amount of serum was added when coating TCPS controls with collagen I. Separately, 15000 EVCMs were suspended in 320 μl of differentiation medium in one well of a low binding 24 well plate. The collagen mix was immediately added to the cells, and the resulting suspension was gently triturated with a 1 ml pipette tip.

### Visualization of t-tubules

EVCMs plated on PAAm gels were incubated with 5μM Di-8-ANEPPS for 10 minutes. The dye was diluted in the following buffer: Phosphate buffered saline with HEPES, 23 mM; glucose, 21 mM; creatine, 5mM; MgCl2, 5 mM. After 10 minutes the dye containing buffer was replaced with the buffer alone and the cells were imaged on a confocal microscope (Nikon A1R). CMs from adult mice were used as a positive control.

### Statistical analysis

Statistical analysis was performed using the two-sided t-test or ANOVA, as described in the figure legends. Differences were considered significant if p was lower than 0.05 (p < 0.05). Error bars represent standard deviations in all figures.

Calculation of myocyte force is described in the Supplementary Information.

## Acknowledgements

We would like to thank the CRM flow cytometry core at MGH for assistance with cell sorting and Dr. Jan Willem Buikema for comments on the manuscript. This work was supported by the Cardiovascular Research Center at the Massachusetts General Hospital.

## Competing financial interests

The author(s) declare no competing financial interests.

